# Inducible biosynthesis of bacterial cellulose in recombinant *Enterobacter* sp. FY-07

**DOI:** 10.1101/2024.06.03.597270

**Authors:** Jiaxun Ren, Liangtian Miao, Wei Feng, Ting Ma, Huifeng Jiang

## Abstract

Bacterial cellulose (BC) is an extracellular polysaccharide with myriad unique properties, such as high purity, water-holding capacity and biocompatibility, making it attractive in materials science. However, genetic engineering techniques for BC-producing microorganisms are rare. Herein, the electroporation-based gene transformation and the λ Red-mediated gene knockout method with a nearly 100% recombination efficiency were established in the fast-growing and BC hyperproducer *Enterobacter* sp. FY-07. This genetic manipulation toolkit was validated by inactivating the protein subunit BcsA in the cellulose synthase complex. Subsequently, the inducible BC-producing strains from glycerol were constructed through inducible expression of the key gene *fbp* in the gluconeogenesis pathway, which recovered more than 80% of the BC production. Finally, the BC properties analysis results indicated that the induced-synthesized BC pellicles were looser, more porous and reduced crystallinity, which could further broaden the application prospects of BC. To our best knowledge, this is the first attempt to construct the completely inducible BC-producing strains. Our work paves the way for increasing BC productivity by metabolic engineering and broadens the available fabrication methods for BC-based advanced functional materials.

## 1. Introduction

Bacterial cellulose (BC) is an extracellular polysaccharide constituting linear chains of glucose residues covalently linked by β-1,4-glycosidic bonds [1]. Compared to the well-known plant cellulose (PC), BC possesses various unique properties such as high crystallinity, ultra purity, hyper water-holding capacity and enhanced biocompatibility [2]. In addition, the productivity of BC (79.2 t/22 days in a 500 m^3^ bioreactor) is significantly higher than that of PC (80 t/7 years in 1 ha of eucalyptus) [3]. Even though the algae have a relatively higher growth rate, the cellulose productivity of algae (less than 36 t/year/ha) cannot be compared with that of cellulose-producing bacteria, either [4, 5]. Due to these superior properties, BC is widely used in food [6], cosmetics [7], biomedical materials [8, 9], lithium battery separators [10] and other areas.

BC can be synthesized by several bacterial genera, including *Gluconacetobacter*, *Azotobacter*, *Pseudomonas*, *Agrobacterium*, *Agrobacterium*, *Salmonella* and *Enterobacter* [2, 11]. Among these BC-producing strains, the species *Gluconacetobacter xylinus* (also known as *Acetobacter xylinus*, or *Komagataeibacter xylinus*) has the remarkable ability to produce BC on a commercial scale [2, 12]. Since *G. xylinus* is strictly aerobic, membranous BC could only form on the gas-liquid interface during the static culture method [13]. Thus, the insufficient oxygen supply during static fermentation results in a long BC production cycle (usually 5∼25 days) [2, 8]. Several genetically modified tools, such as the λ Red and FLP/FRT-mediated site-specific recombination system [14], or the CRISPR interference-mediated gene downregulation method [15] have been reported in *K. xylinus*. Nevertheless, the low growth rate of acetic acid bacteria caused by incomplete glycolysis [10] greatly hampers the improvement of BC production through metabolic engineering. Most efforts in BC researches were focused on the isolation of novel strains [11], the optimization of media composition and culture conditions [12], and BC modifications [2].

The facultative anaerobic BC-producing strain *Enterobacter* sp. FY-07 (FY-07) was isolated from oilfield-produced water to solve the problems encountered by *G. xylinus* [16]. This strain could sharply reduce the BC production cycle and increase BC productivity because of its rapid growth rate. BC synthesis mechanism of FY-07 was studied in the later researches [17]. The BC productivity of FY-07 was enhanced to 13.96 g/L/day by the addition of xanthan gum and KNO_3_ [18, 19]. The water-holding capacity of the BC hydrogel increased by more than 1.7-fold after the overexpression of the colanic acid biosynthetic gene cluster [20]. Despite these marvelous improvements in BC production or modification, genetic engineering of FY-07 still relies on the traditional conjugation transfer combined with the complicated single-crossover and double-crossover selection methods [17]. The most frequently used electroporation-based gene transformation method has already been successfully applied in the cellulose-producing strain *Enterobacter* sp. CJF-002 [21]. Consequently, it is feasible to modify the existing efficient gene transformation methods and genetic manipulation tools for FY-07, paving the way for metabolic engineering of FY-07 to increase BC production.

Almost all the BC producers synthesize BC constitutively. It is still challenging to construct an inducible BC-producing strain as all the genes in the BC biosynthesis pathway are crucial to the normal metabolism of the host [22, 23]. We hypothesized that the inducible BC synthesis strain from glycerol could be constructed through inducible expression of the key gene in the gluconeogenesis pathway (Fig. 1). In this study, the electroporation-based gene transformation and λ Red-mediated gene knockout method were developed for the genetic engineering of FY-07. This toolkit facilitated the construction of the inducible BC-producing strains by controlling the expression of the *fbp* gene in BC biosynthesis pathway (Fig. 1). Our work demonstrated that BC synthesis could be controlled *in vivo* and thus expanded the *in situ* fabrication methods of advanced materials based on BC.

**Fig. 1.**
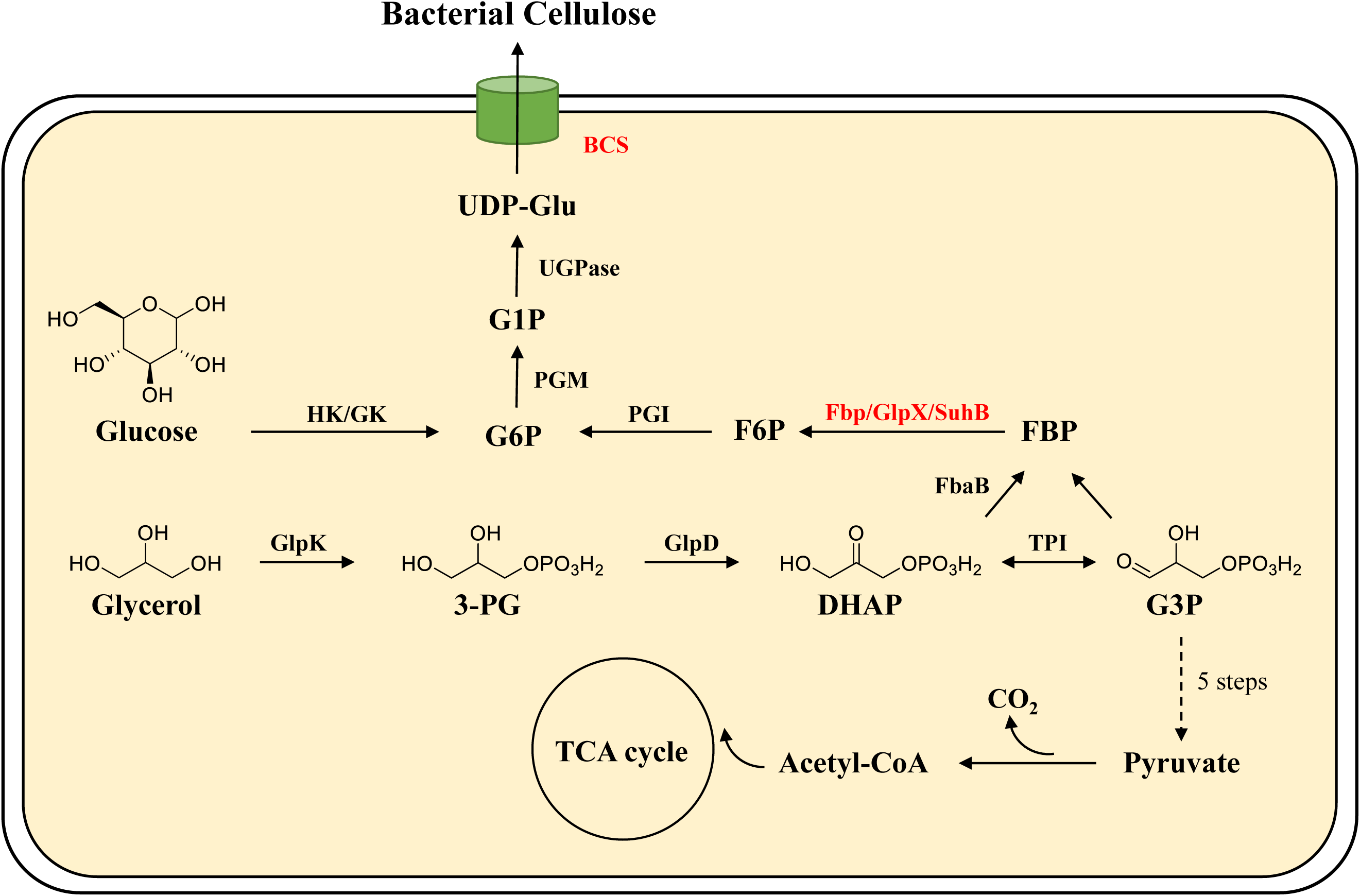
The BC biosynthesis pathway in *Enterobacter* sp. FY-07 using glucose or glycerol as carbon source. Abbreviations: HK: hexokinase; GK: glucokinase; PGM: phosphoglucomutase; UGPase: UDP-glucose pyrophosphorylase; PGI: phosphoglucoisomerase; Fbp: fructose-1,6-bisphosphatase I; GlpX: fructose-1,6-bisphosphatase II; SuhB: inositol-phosphate phosphatase; FbaB: fructose-bisphosphate aldolase class I; TPI: triose-phosphate isomerase; GlpD: aerobic glycerol 3-phosphate dehydrogenase; GlpK: glycerol kinase; G6P: glucose-6-phosphate; G1P: glucose-1-phosphate; UDP-Glu: UDP-glucose; F6P: fructose-6-phosphate; FBP: fructose-1,6-bisphosphate; G3P: glyceraldehyde-3-phosphate; DHAP: Dihydroxyacetone phosphate; 3-PG: 3-phosphate-glycerol.

## 2. Materials and methods

### 2.1. Strains, plasmids, media and culture conditions

The primers, strains and plasmids used in this study are listed in Table 1 and Table 2. *Enterobacter* sp. FY-07 (CGMCC No. 6103), isolated from the production water of the Jilin oilfield (China Petroleum Natural Gas Co., Ltd., Jilin Oil Field Branch), was used as the starting strain for the genetic control of BC biosynthesis [16]. For *Escherichia coli* strain construction, the cells were cultured at 37 °C, 220 rpm in Luria-Bertani (LB) broth (10 g/L tryptone, 5 g/L yeast extract and 10 g/L NaCl). For the construction of FY-07 derivatives, LB broth with 1 g/L cellulase (Celluclast^®^ 1.5 L, Novozymes) was used, and the cells were cultured aerobically at 30 °C. When necessary, appropriate antibiotic (kanamycin 50 mg/L, chloramphenicol 34 mg/L, or spectinomycin 50 mg/L) was added into the medium. For the detection of fluorescence intensity, the refreshed strains were inoculated into 1.8 mL LB broth with cellulase in MicroScreen 48-well cell culture plates at an initial OD_600_ of 0.05 and cultured at 30 °C, 500 rpm for 12 h.

**Table 1.**
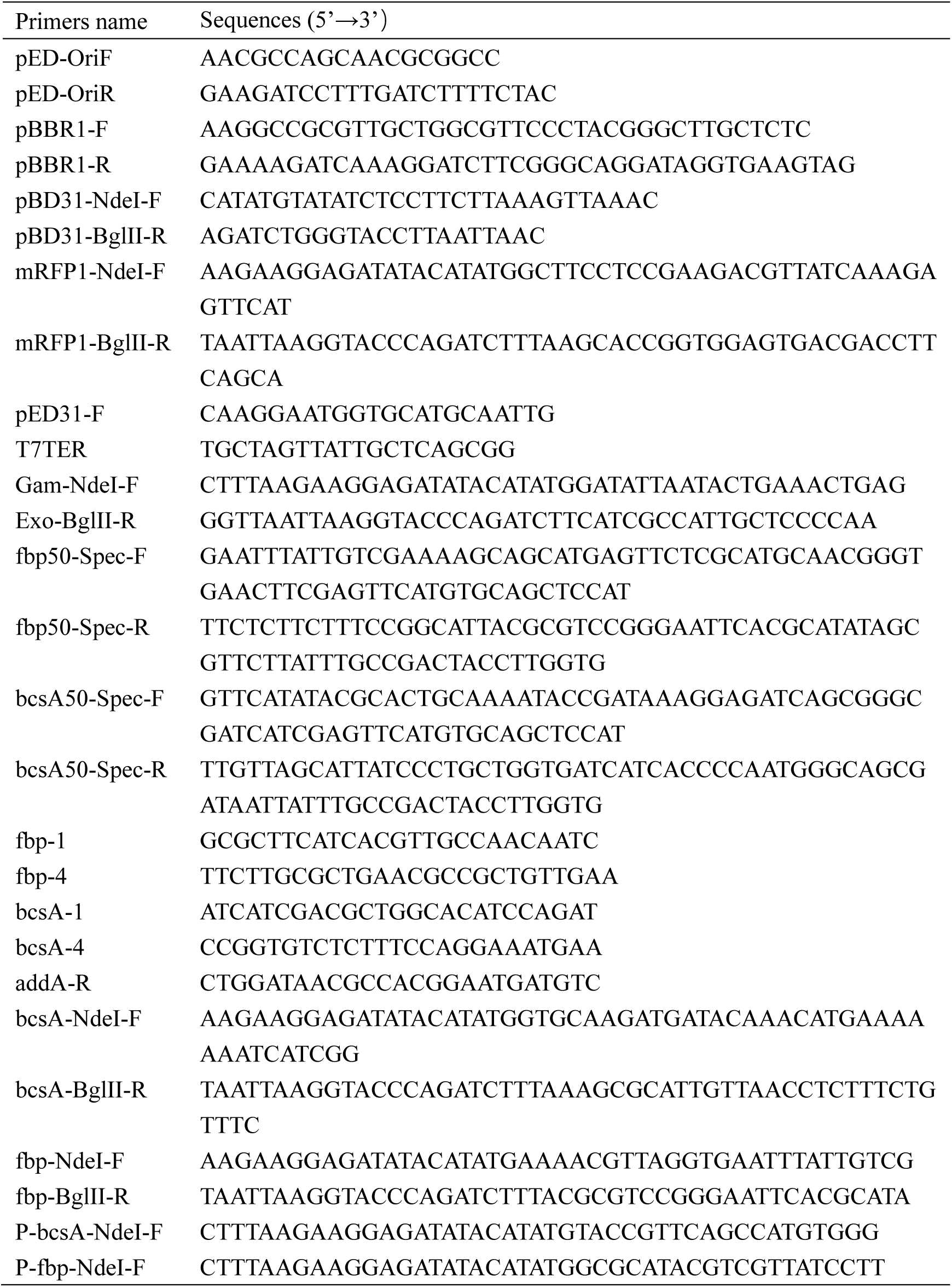
Primers used in this study.

**Table 2.**
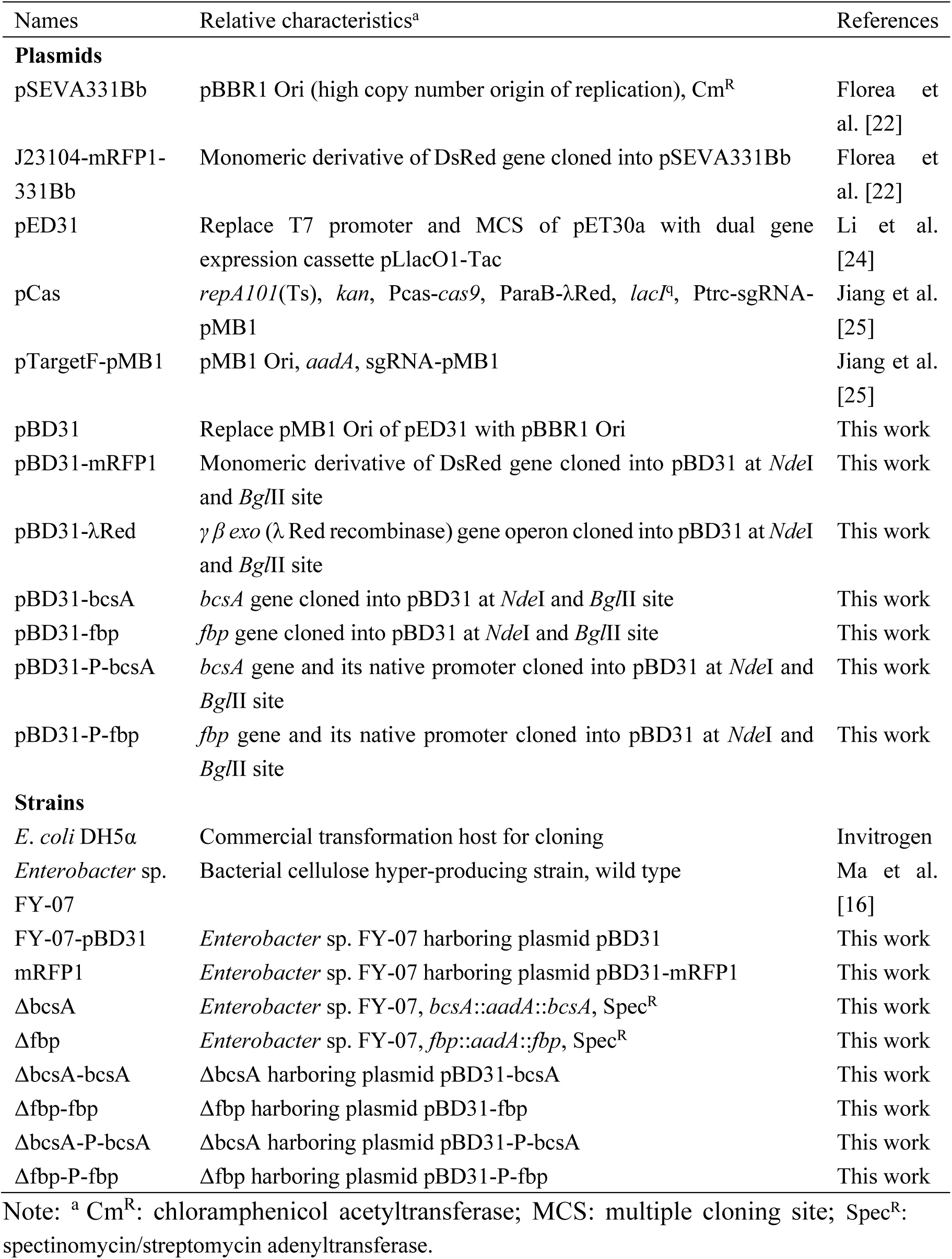
Plasmids and strains constructed in this study.

### 2.2. Construction of recombinant plasmids

The pMB1 replication origin of pED31 [24] was replaced with pBBR1 Ori and pBBR1 Rep protein from pSEVA331Bb vector [22] using a Gibson assembly strategy to yield the expression vector pBD31. The gene encoding a monomeric red fluorescent protein was amplified via PCR from J23104-mRFP1-331Bb [22] with the primer pair mRFP1-NdeI-F/mRFP1-BglII-R, which was then ligated into *Nde*I and *Bgl*II sites of pBD31 to generate pBD31-mRFP1. Similarly, the *γ*, *β* and *exo* gene operon encoding λ Red recombinase from pCas [25] was inserted between *Nde*I and *Bgl*II sites of pBD31 to create pBD31-λRed. The genes encoding fructose-1,6-bisphosphatase I (*fbp*, GenBank: AMO50847.1) and the catalytic subunit of cellulose synthase (*bcsA*, GenBank: AMO47317.1) in FY-07 were cloned into pBD31 using the same strategy to yield the expression vectors pBD31-fbp and pBD31-bcsA, respectively. The native promoters of *fbp* (285 bp) and *bcsA* (171 bp) were also inserted into pBD31-fbp and pBD31-bcsA separately so as to enhanced their expression levels, resulting in the plasmids pBD31-P-fbp and pBD31-P-bcsA.

### 2.3. Construction of recombinant strains

The electroporation-based gene transformation method for *E. coli* [26] was modified for genetic engineering in FY-07. Briefly, newly refreshed FY-07 cells were cultured in 30 mL LB broth with cellulase at 30 °C, 220 rpm to an OD_600_ of 0.6∼0.8 and then standing statically for 2 h at room temperature until all the BC was decomposed by cellulase. These cells were made electrocompetent by concentrating 100-fold and washing four times with ice-cold 10% glycerol. Electroporation was performed according to the manufacturer’s instructions by using 50 μL of cells and 100 ng of plasmids/purified PCR products. Shocked cells were added to 1 mL LB with cellulase, incubated for 4 h at 30 °C, and then spread onto appropriate agar plate to select the right transformants.

A one-step homologous recombination method [27] was applied to integrate the *aadA* gene encoding spectinomycin/streptomycin adenyltransferase into the FY-07 chromosome at the *bcsA* or *fbp* site, inactivating the *bcsA* or *fbp* gene. Taking the *bcsA* gene deletion as an example, the *aadA* gene was amplified from pTargetF-pMB1 using the primer set bcsA50-Spec-F/bcsA50-Spec-R, which was digested by *Dpn*I to remove the plasmid template and electroporated into the competent cell of FY-07 harboring pBD31-λRed. After overnight growth on LB plate with kanamycin and spectinomycin, several colonies were picked for PCR verification and sequencing using the primer set bcsA-1/bcsA-4. The helper plasmid pBD31-λRed was cured by serial growing the right colonies overnight in the medium without kanamycin. After determining their sensitivity to kanamycin, the right colony was designated ΔbcsA. The *fbp* gene was knocked out of the FY-07 chromosome via the same method, yielding the recombinant strain Δfbp.

### 2.4. BC production and purification

The BC-producing strains were precultured in 5 mL LB medium with cellulase for 24 h. This seed culture was inoculated into 50 mL modified Hestrin-Schramm (mHS) medium [19] (2.5% glycerol, 1% tryptone, 0.75% yeast extract and 1% disodium phosphate anhydrous) in 250 mL flasks at an initial OD_600_ of 0.05 and cultured statically at 30 °C. The obtained BC films were treated with 0.1 M NaOH at 100 °C for 30 min to remove the attached cells and media. The BC was washed with deionized water until it turned white, and the pH of the BC became neutral. Then, the purified BC was dried at 60 °C for overnight or freeze-dried until it reached a stable weight and weighed.

### 2.5. Analytical methods

Cell growth was analyzed by measuring the optical density at 600 nm using a V-1800 spectrophotometer (Mapada, China). The fluorescence intensity of the monomeric red fluorescent protein (mRFP1) was measured with a BioTek multi-mode microplate reader (Synergy Neo2, USA; excitation 590 nm, emission 630 nm) [14]. The expression of mRFP1 in the cells was further analyzed via a laser scanning confocal microscopy (Leica TCS SP5 II, Germany).

### 2.6. BC properties analysis methods

#### 2.6.1. Scanning electron microscopy (SEM)

For the bacteria and cellulose complex analysis, BC pellicles formed after 10 days’ cultivation were cut into small pieces, washed with phosphate buffered saline and fixed with 1% glutaraldehyde and 0.4% paraformaldehyde. Then, the fixed samples were dehydrated in a graded series of water-ethanol solutions ranging from 30% to 100% ethanol and dried by a critical point dryer (Leica EM CPD030). The top surface of the BC pellicle was coated with platinum under vacuum and observed via a scanning electron microscopy (Hitachi SU8010, Japan) operating at 1.0 kV [22, 28].

For the BC material analysis, the freeze-dried BC membranes were cut into small pieces, mounted on aluminum studs, and coated with platinum under high-vacuum conditions. The microstructures of the BC membranes were observed via SEM operating at 3.0 kV [20, 29].

#### 2.6.2. Fourier transform infrared spectroscopy (FT-IR)

The infrared spectra of the freeze-dried BC samples were measured using a Nicolet iS5 Fourier transform infrared spectrometer (Thermo Scientific, America). All of the spectra were recorded with a resolution of 2 cm^−1^ ranging from 4000 to 500 cm^−1^ [16, 30].

#### 2.6.3. X-ray diffraction (XRD)

The crystallinity of the freeze-dried BC samples was determined by a SmartLab X-ray diffractometer (Rigaku Corporation, Japan) with Ni-filtered CuK (k=1.54 Å) radiation. Scans were performed over the 5-80°, speed of 10°/min, and 2θ range using steps of either 0.02° in width. The degree of crystallinity was calculated according to the formula: CrI(%)=(1−(I_AM_/I_200_))×100% [30, 31].

### 2.7. Statistical analysis

All experiments were repeated at least three times. The significance of the data was evaluated using independent-samples *t*-test with *P*<0.05 indicating significance.

## 3. Results

### 3.1. Development of a gene overexpression vector for *Enterobacter* sp. FY-07

Since the genome of FY-07 is highly similar to that of *E. coli* [17], we hypothesized that the universal genetic engineering toolkits could be amended for FY-07. We used the well-characterized plasmid pED31 for *E. coli* [24] as the backbone to create a gene overexpression vector for FY-07. Firstly, the pMB1 replication origin of pED31 was replaced with the replication origin and its Rep protein for replication of the broad-host-range plasmid pBBR1 from *Bordetella bronchiseptica* [22], which has been demonstrated to replicate in *Acetobacteraceae* [14]. Next, the monomeric red fluorescent protein (mRFP1) was used as a reporter protein to determine whether pBD31 could replicate in FY-07 (Fig. 2A). This recombinant plasmid was then used to develop protocols for the preparation of electrocompetent cells and DNA transformation of FY-07 (see 2.3.). The recombinant strain mRFP1 showed visually detectable expression of red fluorescent protein after overnight induction with 1 mM isopropylthiogalactoside (IPTG). Almost all of the cells emitted clear cytosolic red fluorescence under the laser scanning confocal microscopy (Fig. 2B). In parallel, six different IPTG concentrations (0.05, 0.10, 0.20, 0.30, 0.40, and 0.50 mM) were applied to determine the optimal inducer concentration. Result suggested that cells produced the highest fluorescence intensity per unit OD with 0.4 and 0.5 mM IPTG (Fig. 2C). In addition, the BC pellicle synthesized by the mRFP1 strain was also stained with visual red fluorescence after static fermentation (Fig. 2D). These results indicated that the pBBR1 Ori replicator and the inducible promoter P_LacO1_-lacI^q^ can be suitable for controlling gene expression in FY-07.

**Fig. 2.**
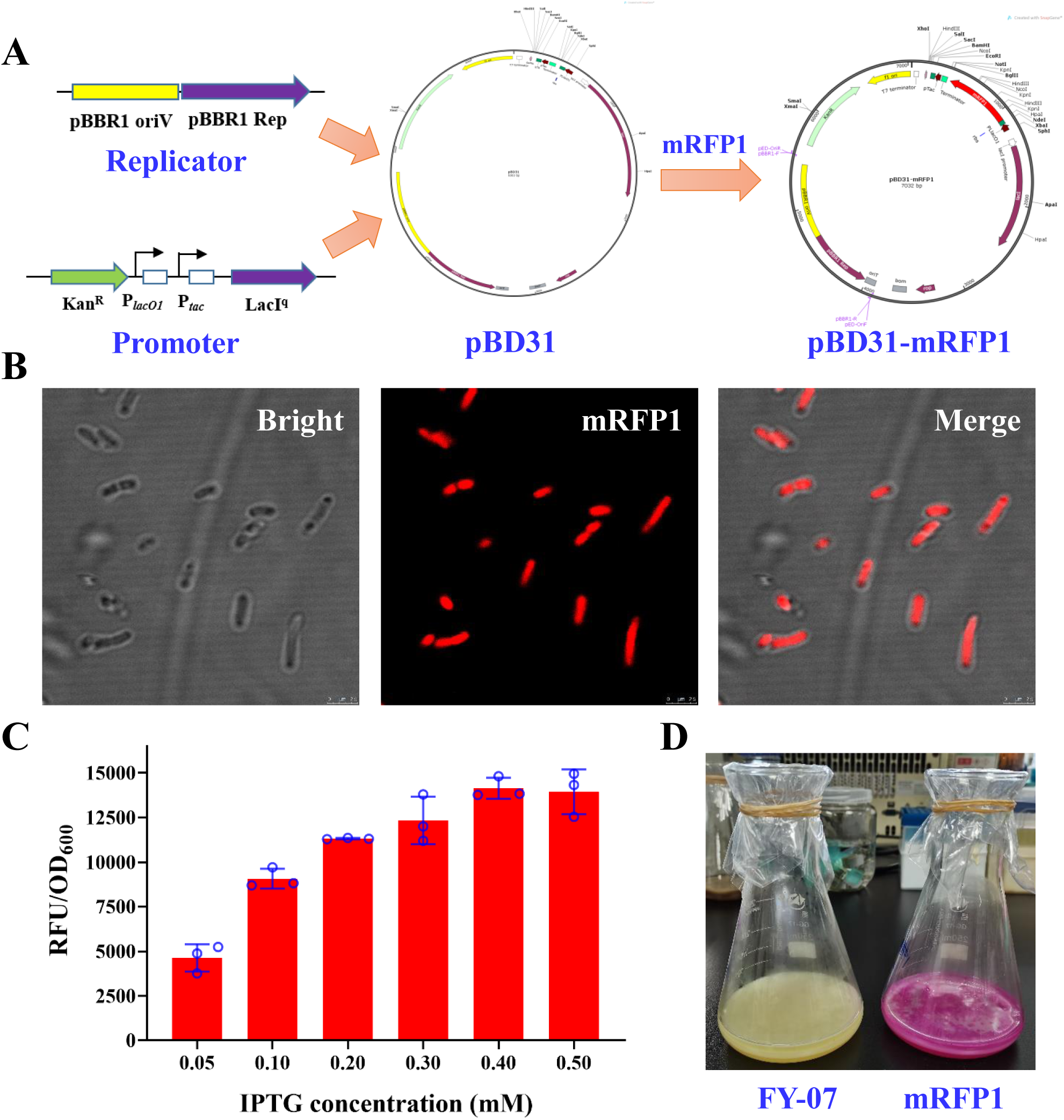
Construction and characterization of the inducible vector pBD31 for *Enterobacter* sp. FY-07. A. Construction workflow of pBD31-mRFP1; B. Laser scanning confocal microscopy images of FY-07 expressed mRFP1. Scale bar, 2.5 μm; C. Comparison of the fluorescence of the mRFP1 strain supplemented with different concentrations of IPTG; D. Morphology of the BC pellicles synthesized by the wild-type (FY-07) and mRFP1.

### 3.2. Development of a λ Red-mediated gene knockout method for FY-07

As BcsA and BcsB form the catalytically active core of BC synthase, inactivating BcsA led to the insufficient BC synthesis [17, 32]. We selected *bcsA* as a target gene to develop the gene knockout method for FY-07. The simple and highly efficient λ Red and FLP/FRT-mediated site-specific recombination system was widely used in various bacteria like *E. coli* [26]. However, we failed to apply this system to delete genes in FY-07 (Fig. S1A) presumably due to the poor expression of λ Red recombinase (Fig. S1B). Therefore, the mRFP1 encoding gene in pBD31-mRFP1 was substituted by *γ*, *β* and *exo* gene operon to improve λ Red expression (Fig. 3A). Homologous recombination events occurred in FY-07 after IPTG-induced λ Red recombinase expression and electroporation of a DNA fragment consisting of a spectinomycin selection marker and 50-nt flanking homologous arms (Fig 3B). Colony PCR verification results showed the different bands of *bcsA* gene between the wild type strain (Lane C, ∼3.0 kb) and ΔbcsA strain (Lanes 1-3, ∼2.0 kb) (Fig. 3C). Further sequencing data also confirmed the deletion of the *bcsA* gene.

**Fig. 3.**
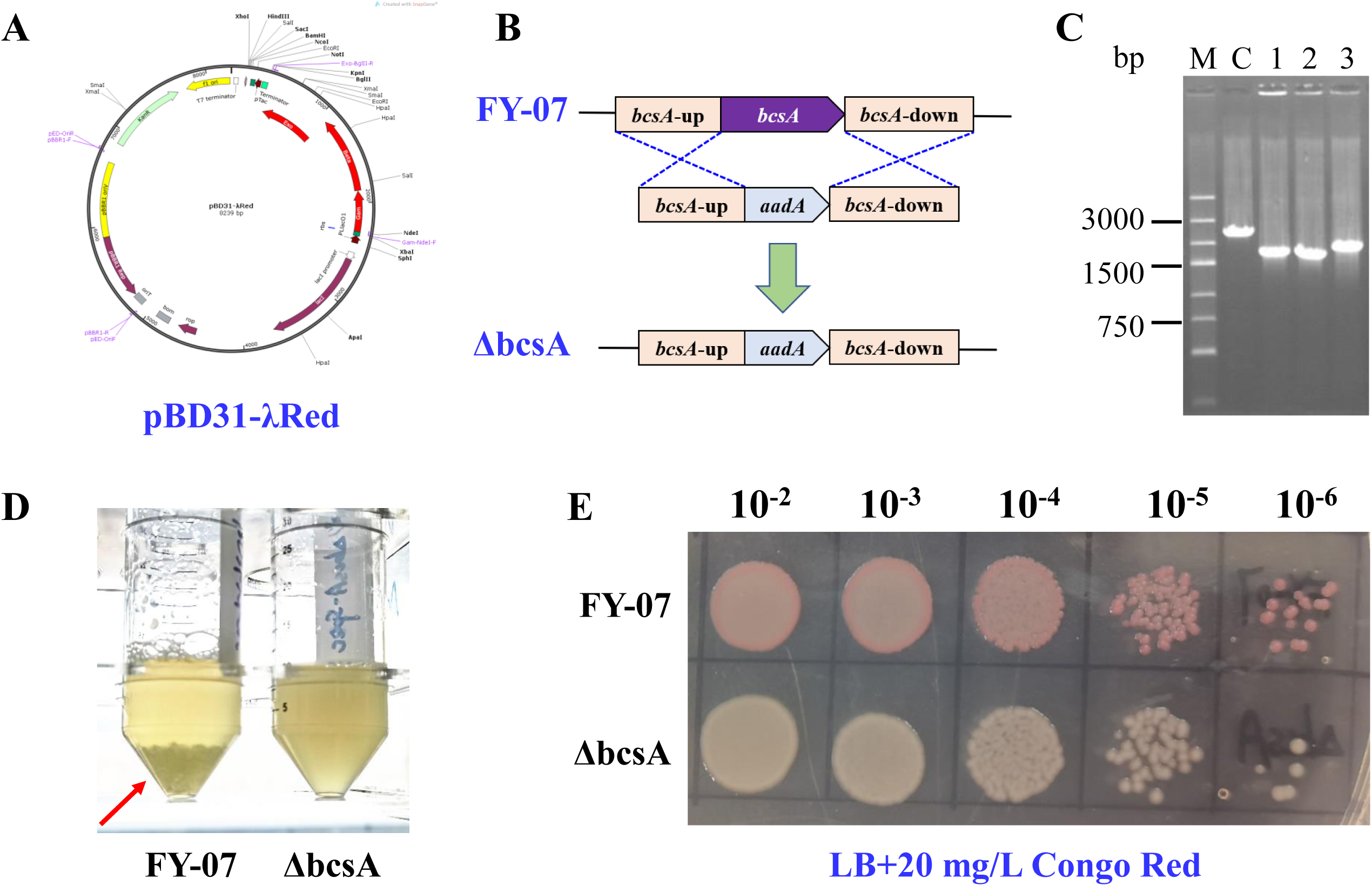
Development of the λ Red-mediated gene knockout method for FY-07. A. Plasmid map of pBD31-λRed; B. Schematic diagrams of the deletion of the *bcsA* gene; C. Colony PCR verification of the ΔbcsA strain (M: marker; C: control); D. Morphology differences between FY-07 and ΔbcsA under agitated culture; E. Effects of *bcsA* gene inactivation on cell growth and BC production.

Similar to *Acetobacteraceae*, the wild type FY-07 produced sphere-like or amorphous BC in agitated culture, which was easily precipitated in the liquid media. This phenomenon disappeared after knocking out *bcsA* gene (Fig. 3D). The cell growth was compared by spotting these two strains onto LB plate supplemented with 20 mg/L Congo Red, which has a strong affinity for cellulose and usually used to evaluate the BC synthesis capacity of different strains [17, 19]. There was no significant impact on cell growth followed by the inactivation of the *bcsA* gene (Fig. 3E). These results suggested the successful development of the gene knockout method for FY-07.

### 3.3. Construction and evaluation of the inducible BC-producing strains

It is desirable to construct inducible BC-producing strains as constitutive cellulose synthesis complicates genetic engineering techniques, and the high metabolic burden caused by cellulose production might lead to cellulose-nonproducing mutants [22]. Nevertheless, inducible expression of *bcsA* in the ΔbcsA strain failed to restore BC synthesis capacity (Fig. S2), probably because of the difficulty in correctly expressing the membrane-anchored protein BcsA in FY-07. The transmembrane region of BcsA was predicted by an online tool (https://www.novopro.cn/tools/tmhmm.html) [33], suggesting eight transmembrane regions in BcsA (Fig. S3). We speculated that inactivation and inducible expression of other key genes involved in the BC biosynthesis pathway like glucokinase, phosphoglucomutase, or UDP-glucose pyrophosphorylase (UGPase) might cause immense problems as these genes affected the normal metabolism of FY-07 [17]. A feasible approach to construct the inducible BC-producing strains is thus to conditionally supply the substrate (glucose) concentration in the BC pathway of FY-07.

Fructose-1,6-bisphosphatase I (Fbp) catalyzes the irreversible conversion of fructose-1,6-bisphosphate to fructose-6-phosphate in the gluconeogenesis pathway (Fig. 4A). A previous study suggested that the *fbp* (AKI40_4472) knockout mutant could not synthesize BC when glycerol was used as the sole carbon source, but the BC production ability remained unaffected in HS medium [23]. According to these references, *fbp* gene was selected as the target gene to construct the inducible BC-producing strains. We knocked out the *fbp* gene via the aforementioned λ Red-mediated gene knockout method. SEM images confirmed that the *fbp* deletion mutant lost its ability to synthesize BC during static culture in mHS medium (Fig. 4B). The genetic replenishment of *fbp* using the P_LacO1_-lacI^q^ promoter or its native promoter (285 bp) was carried out to rescue the BC production deficiency in FY-07. The BC synthesis ability was restored after IPTG induction (Fig. 4B, 4D). However, the fibers of the BC pellicles synthesized by the recombinant strains were not dense compared to those of the wild type according to the SEM images (Fig. 4B). Further SDS-PAGE result revealed that the Fbp (36.94 kDa) expression level under the control of the P_LacO1_-lacI^q^ promoter was markedly higher (Fig. 4C, lane 3) than that under the control of the native promoter (Lane 4) or the wild type (Lane 1). After 7 days’ induced cultivation, the BC yield of these strains reached 6.61, 5.36 and 3.72 g/L, respectively. Notably, the BC production by the recombinant strains (Δfbp-fbp and Δfbp-P-fbp) recovered about 81% (*P*=0.092) and 56% (*P*=0.008), respectively (Fig. 4D). In addition, both strains encountered a leakage expression of Fbp when these inducible promoters were used (Fig. 4D).

**Fig. 4.**
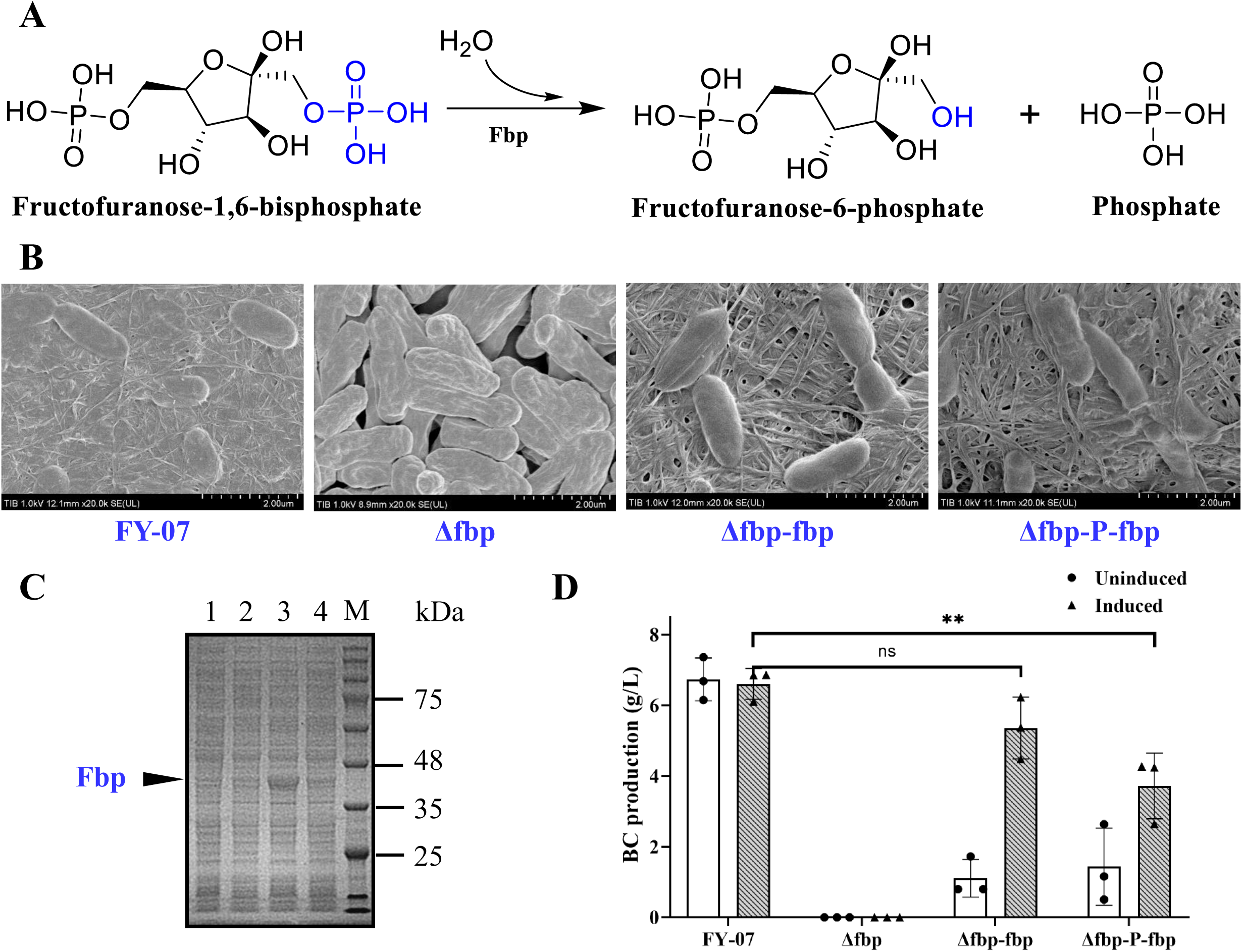
Construction and evaluation of the inducible BC-producing strains. A. Reactions catalyzed by fructose-1,6-bisphosphatase I; B. SEM images of the wild-type and recombinant strains under static condition. Scale bar, 2.0 μm; C. SDS-PAGE profile of the expression of Fbp by different strains (Lanes 1-4: FY-07, Δfbp, Δfbp-fbp, and Δfbp-P-fbp); D. BC production by different strains under static condition. ns: *P*>0.05; *: *P*<0.05; **: *P*<0.01.

### 3.4. BC properties analysis

The microstructures of the BC pellicles synthesized by the wild type and recombinant strains were observed via SEM. All the BC samples formed a reticulated structure consisting of cellulose nanofibrils. The BC synthesized by the wild type strain displayed a denser network structure, while a loose and porous structure was seen in both of the BC pellicles synthesized by the recombinant strains (Fig. 5A). These differences in network structure were consistent with the above-mentioned microscopy images of the bacteria and cellulose complex (Fig. 4B). The change of BC reticular structure resulted in the reduced tension strength and Young’s modulus (Table S2). The storage modulus (G’) and loss modulus (G”) of the induced-synthesized BC samples were also decreased compared to those of the wild type (Fig. S4). Several studies have suggested that the network structure of BC is closely related to its crystallinity [29, 34, 35]. Therefore, FT-IR and XRD were used to obtain further information on the differences between the constitutively synthesized and induced synthesized BC samples.

**Fig. 5.**
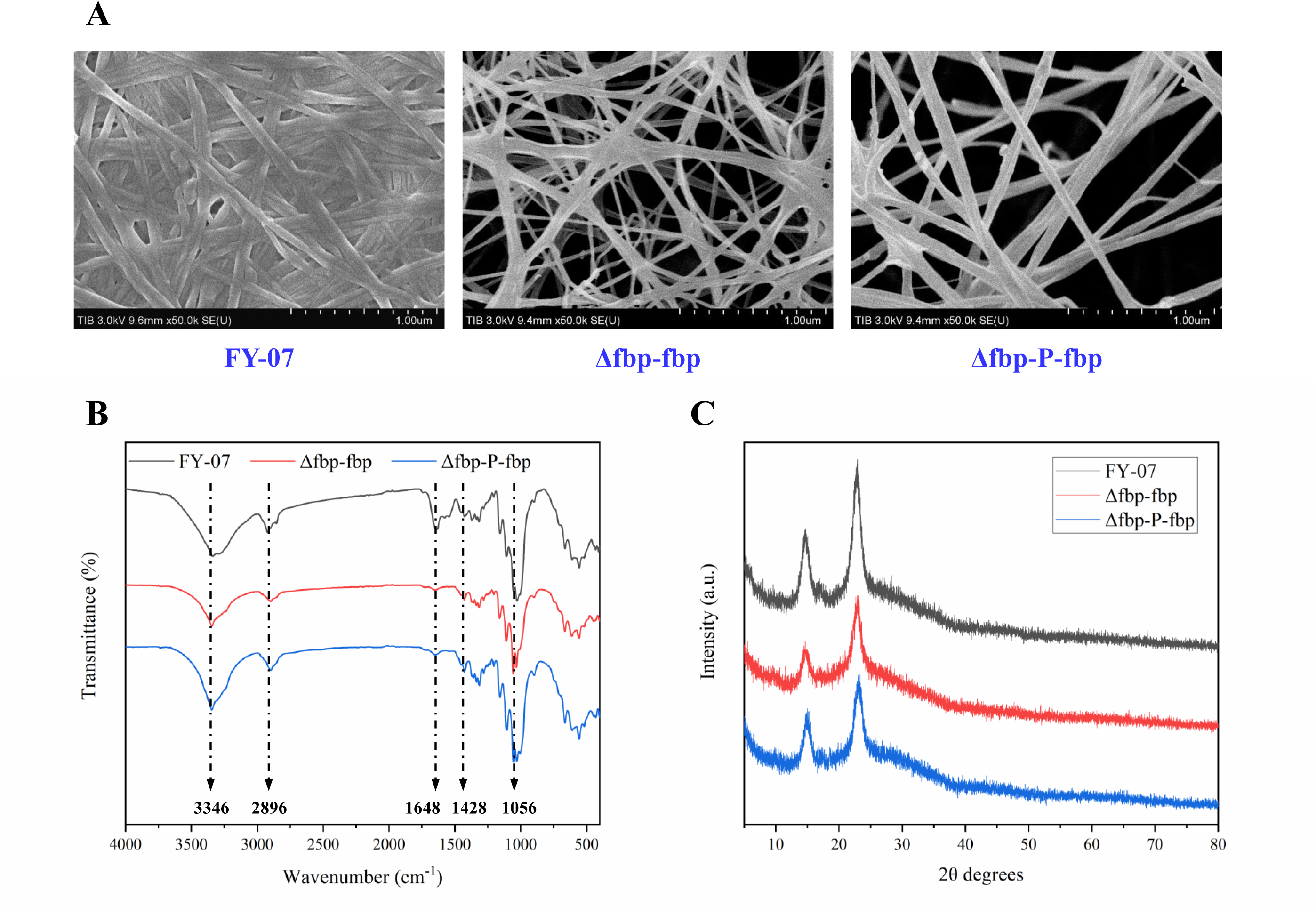
The structural properties of BC produced by the wild-type and recombinant strains. SEM images (A), FT-IR spectra (B) and XRD patterns (C) of the constitutively and induced synthesized BC samples. Scale bar, 1.0 μm.

Preliminary characterization of the BC samples synthesized by the constitutively and inducible BC-producing strains was carried out using FT-IR spectroscopy. The FT-IR spectra of BC pellicles synthesized by different strains matched quite closely (Fig. 5B). The strong absorption peak at 3346 cm^−1^ was the stretching vibration absorption peak of O─H, while the peak at 2896 cm^−1^ corresponded to the stretching vibration absorption peak of C─H, which confirmed the presence of amorphous cellulose [36, 37]. The absorption peaks near 1648, 1428 and 1056 cm^−1^ could be assigned to the carbonyl groups (C═O), methylene (─CH_2_) and C─O─C stretching, respectively (Fig. 5B). The spectra of all the BC samples were basically consistent with those of previous studies, which displayed the typical peaks of cellulose in the FT-IR spectra [18, 28, 29]. The microstructural changes and crystallinity variations in the BC samples produced by these strains were compared through XRD analysis. Similar to the FT-IR results, all the XRD patterns showed the representative diffraction peaks of BC and also matched closely (Fig. 5C). Two dominant diffraction peaks at approximately 14.7° and 22.9° for all the BC samples represented both the crystalline phases Iα and Iβ [28, 30, 38], indicating that the induced BC production mode had no significant effect on the crystal structure of the BC. The CrI values of the constitutively synthesized and induced synthesized BC samples calculated from the XRD diffractograms were 71.43%, 61.54% and 59.89%, respectively. The reduction in crystallinity under inducible BC synthesis conditions may be the result of changes in the arrangement of BC chains during BC production [29, 39].

## 4. Discussion

It is attractive to construct inducible BC producing strains as the separation of cell growth with BC synthesis would avoid cellulose-nonproducing mutants [22]. The inducible BC synthesis facilitates *in situ* fabrication of functional BC materials with a low degree of crystallinity (Fig. 5C). This low-crystallinity BC shares diverse advantages especially in biomedical or cosmetics areas [40]. For example, BC with low crystallinity results in high rehydration and water holding capacity, which helps to absorb tissue fluid and blood exuded from the wound to achieve hemostasis [41]. The proper water holding capacity of dressing materials has proved to accelerate the wound healing process [42]. In addition, low crystallinity BC shows improved dispersibility in water, which is useful as excipients in preparation of tablets and confectionery products [43]. Therefore, reducing the crystallinity can widen the application prospects of BC. Several attempts have been made to engineer the control of BC production. Florea et al. [22] reported that BC production could be suppressed by inhibiting UGPase translation via synthetic regulatory small RNA (sRNA) technology. Recently, a conditional BC-producing strain was constructed by Gao et al. [23] through deletion of the *fbp* gene. This strain could produce BC only using glucose rather than glycerol as the sole carbon source [23]. Unfortunately, all of those studies failed to obtain completely inducible BC-producing strains. In this research, an inducible BC-producing strain was generated by controlling Fbp expression with the inducible promoter P_LacO1_-lacI^q^, which is the first demonstration of the separation of cell growth with BC synthesis. FT-IR, thermogravimetric analysis (TGA) and differential scanning calorimetry (DSC) results indicated that most BC properties of the recombinant strains were similar to those of the wild type (Fig. 5, S5). More importantly, the network structure of BC changed after the *fbp* expression level was altered (Fig. 4C, 5A). The looser and more porous structure of the induced synthesized BC was prone to be the result of reduced BC production yield, as the BC membrane porosity has a negative relationship with the corresponding BC yield (Fig. 4D, 5A). The variability of the BC membrane porosity caused by the different BC yield was consistent with a previous report [15]. Accordingly, the regulation of the BC reticular structure could be achieved by precisely controlling the expression of key genes in the BC biosynthesis pathway, as the ratio of BC yield determine the BC structure. This tunability of the network structure endows BC with unique technological appeal for the fabrication of advanced functional materials like porous membranes [44]. Finally, the convenient and highly efficient electroporation-based gene transformation and λ Red-mediated gene knockout method was applied in the BC-producing strain FY-07, paving the way for metabolic engineering of FY-07.

Genetic engineering is one of the most efficient strategies for increasing the production of target metabolites or for valorizing waste into value-added products [45, 46]. Nevertheless, there are only a few reports on the use of genetic engineering techniques to enhance BC production, mainly because of the lack of genetic toolkits for systematic strain engineering [10, 14, 47]. A genetic toolkit consisting of various plasmids, promoters and reporter proteins has been characterized in *Acetobacteraceae* [22]. This toolkit was expanded by adding and characterizing additional promoters, terminators, ribosome binding sites and degradation tags in a subsequent study [47]. The λ Red-mediated homologous recombination system and CRISPR interference system have been successfully applied in *K*. *xylinus* in recent reports [14, 15]. In our work, a gene overexpression vector and the λ Red-mediated gene knockout method were constructed and characterized for the fast-growing and BC hyperproducing strain FY-07. The overexpression of λ Red recombinase dramatically improved the gene knockout efficiency in FY-07 (Fig. S1). Additionally, one-step inactivation of chromosomal genes in FY-07 using PCR products is more convenient and efficient (with a nearly 100% recombination efficiency, Table S1) than the traditional suicide plasmid-based gene deletion method [21, 26]. However, the cumbersome marker recycling steps cannot be ignored in this system. The scarless CRISPR/Cas9 system is currently under construction for this strain.

Dephosphorylation of fructose-1,6-bisphosphate plays a crucial role in gluconeogenesis when glycerol is used as the sole carbon source [23]. We demonstrated that the expression of Fbp with the IPTG-induced promoter P_LacO1_-lacI^q^ in FY-07 enabled the inducible production of BC pellicles. Actually, BC productivity from glycerol (∼1 g/L/d) is significantly lower than that from glucose (∼14 g/L/d) [19] in FY-07 (Fig. 4D). A similar phenomenon was also found in *Acetobacteraceae* [48]. The presumed reasons mainly include the following: (1) the BC biosynthesis pathway from glycerol (9 steps) is more complicated than that from glucose (4 steps); (2) The substrate transport system for glucose (phosphotransferase system) is much more efficient than the GlpF-aided glycerol transport system [49]; (3) More oxygen is required to synthesize biomass and BC when glycerol is used as the substrate since glycerol is more reduced than glucose [50]. Interestingly, the BC yield of the recombinant strains decreased even though the Fbp expression level in the recombinant strains was much higher than that of the wild type strain (Fig. 4). The reason is that most resources allocated to one specific protein cause metabolic imbalance (or metabolic burden), leading to a decreased target production. This phenomenon has been commonly observed in previous reports [27, 51]. Other efficient inducible gene expression tools, such as optogenetic tools [52, 53] or thermosensitive bio-switches [54], should be exploited for the precise and dynamic regulation of target protein expression in order to achieve higher BC production.

Another noteworthy phenomenon observed in our study was the leakage expression of Fbp using the P_LacO1_-lacI^q^ promoter in FY-07. A small amount of BC was also produced by Δfbp-fbp and Δfbp-P-fbp even without the addition of IPTG (Fig. 4D). In contrast, there was no BC synthesized by Δfbp, illustrating that the production of BC by Δfbp-fbp and Δfbp-P-fbp under uninduced condition was not the result of Fbp isoenzymes. Leakage expression by the P_LacO1_-lacI^q^ promoter was also found in the inducible expression of mRFP1 (data not shown). To solve the leakage expression problem caused by inducible promoters, many researchers have designed orthogonal promoter libraries to screen for extremely low leakage or even zero-leakage expression system for toxic protein production [55–57]. The power of these low leakage expression systems will be harnessed in future studies.

## 5. Conclusion

In this study, the electroporation-based gene transformation and the λ Red-mediated gene knockout method were developed for the first time for *Enterobacter* sp. FY-07. The inducible BC-producing strains from glycerol were constructed through inducible expression of the key gene *fbp* in the gluconeogenesis pathway. To our best knowledge, this is the first attempt to construct the completely inducible BC-producing strains. The properties of the BC pellicles from the recombinant strains and the wild-type strain FY-07 were compared. The SEM results indicated that the induced-synthesized BC pellicles were looser and more porous, which was confirmed by tensile and rheological tests. The change of BC reticular structure was prone to be the results of the reduced BC production yield. The FT-IR results showed that the O─H, C─H, C═O, ─CH_2_ and C─O─C groups were basically consistent with the organic compound groups contained in the molecular structure formula of cellulose synthesized by other BC producers. There were no notable differences among these BC samples according to the TGA and DSC results. A slight reduction in crystallinity also occurred during the inducible BC production process. The reduction in crystallinity might be the result of the reduced changes in the arrangement of BC chains during BC production. This work provides an efficient toolkit for the metabolic engineering of FY-07 and broadens the available fabrication methods for BC-based advanced functional materials.

## CRediT authorship contribution statement

**Jiaxun Ren:** Methodology, Formal analysis, Investigation, Validation; **Liangtian Miao:** Conceptualization, Methodology, Formal analysis, Validation, Writing – original draft; **Wei Feng:** Methodology, Investigation; **Ting Ma:** Writing – review & editing, Supervision; **Huifeng Jiang:** Writing – review & editing, Funding acquisition, Project administration, Supervision.

## Declaration of competing interest

The authors declare that they have no known competing financial interests or personal relationships that could have appeared to influence the work reported in this paper.

## Data availability

Data will be made available on request.

## Acknowledgements

This work was supported by the National Key R&D Program of China (Grant No. 2021YFC2103500). We thank Huanhuan Zhai from the core facility center at Tianjin Institute of Industrial Biotechnology for her technical advice on the SEM experiments.

## Notes

### Competing Interest Statement

The authors have declared no competing interest.

