## Supporting information for "Inducible biosynthesis of bacterial cellulose in recombinant *Enterobacter* sp. FY-07"

### TITLE

---

[§] These authors contributed equally to this work.

[*] Corresponding author.

### Supplementary Materials and Methods

#### *Mechanical properties*

A texture analyzer (DMA Q800, TA Instruments, USA) was used to measure BC mechanical properties according to ASTM D882. The purified BC membranes ere cut into strips (2×7 cm). The initial grip separation and crosshead speed used were set at 40 mm and 1 mm/s, respectively. In each test, Tensile strength (TS) and elongation (E%) were calculated using Eqs. (1) and (2), respectively.

$$TS=F/(L\times X) \quad (1)$$

$$E\%=100\%\times(l-l_1)/l_1 \quad (2)$$

Where $F$ is the tensile force (N), $L$ the width of the film (mm), $X$ the thickness (mm), $l_1$ is the initial length of the film and $l$ is the length of the film at breaking point. The slope of the curve in the linear elastic region was calculated to obtain Young's modulus.

#### *Rheological properties*

The rheological properties of freeze-dried BC were determined in a dynamic rheometer (ARES-G2, TA Instruments, USA) using a small amplitude oscillatory test (SAOT). The dynamic rheometer was equipped with rough surface parallel-plate geometry (20 mm diameter) to prevent sample slippage. Gap and strain were set at 2.0 mm and 1.0%, respectively. Steady shear tests were performed over a shear rate range of 0.065–65 1/s and conducted between 0.01 and 10 Hz at 20 °C in frequency sweeps. Three replicate scans were conducted, with storage modulus (G') and loss modulus (G'') recorded.

***Thermogravimetric analysis (TGA) and differential scanning calorimetry (DSC)***

The TGA and DSC of the freeze-dried BC pellicles were carried out by a TGA/DSC 1/1600 simultaneous thermal analyzer (METTLER TOLEDO, Switzerland) to test its heat resistance. The experimental conditions were 25–800 °C, 10 K/min, and $N_2$ was fed at a flow rate of 30.0 mL/min.

### Supplementary Results

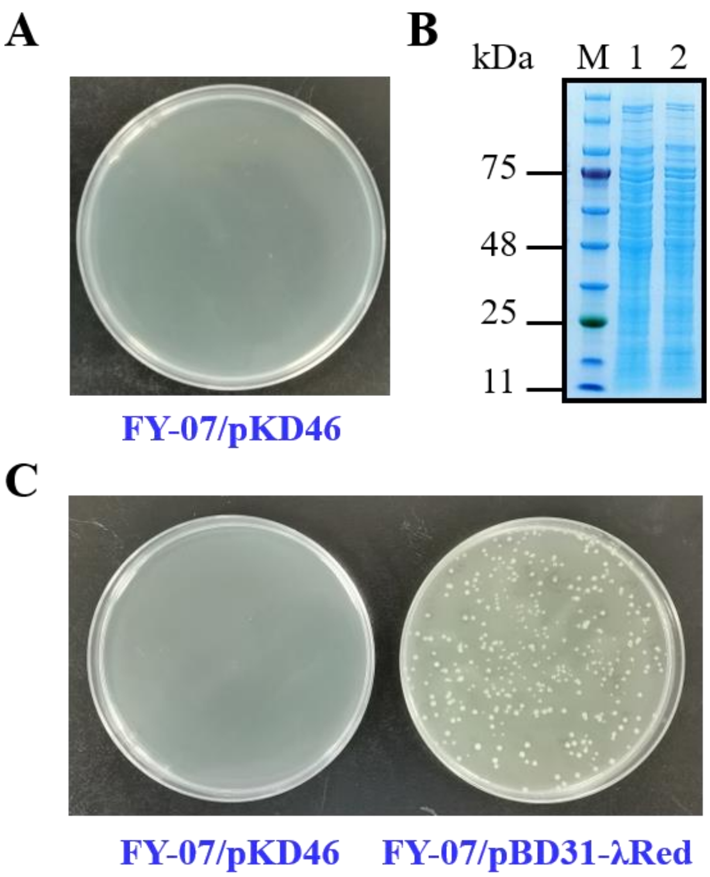


**Fig. S1 Knocking out of *bcsA* in FY-07 chromosome with the helper plasmid pKD46 or pBD31-λRed**

Note: A: The DNA fragments for *bcsA* deletion was electroporated into the competent cell of FY-07 harboring pBD31-λRed; B: SDS-PAGE profile of the expression of λ-Red recombinase subunits Gam (16.3 kDa), Beta (29.7 kDa) and Exo (25.9 kDa) in FY-07/pKD46 (Lanes 1-2: FY-07, FY-07/pKD46); C: The DNA fragments for *bcsA* deletion were electroporated into the competent cell of FY-07 harboring pKD46 or pBD31-λRed.

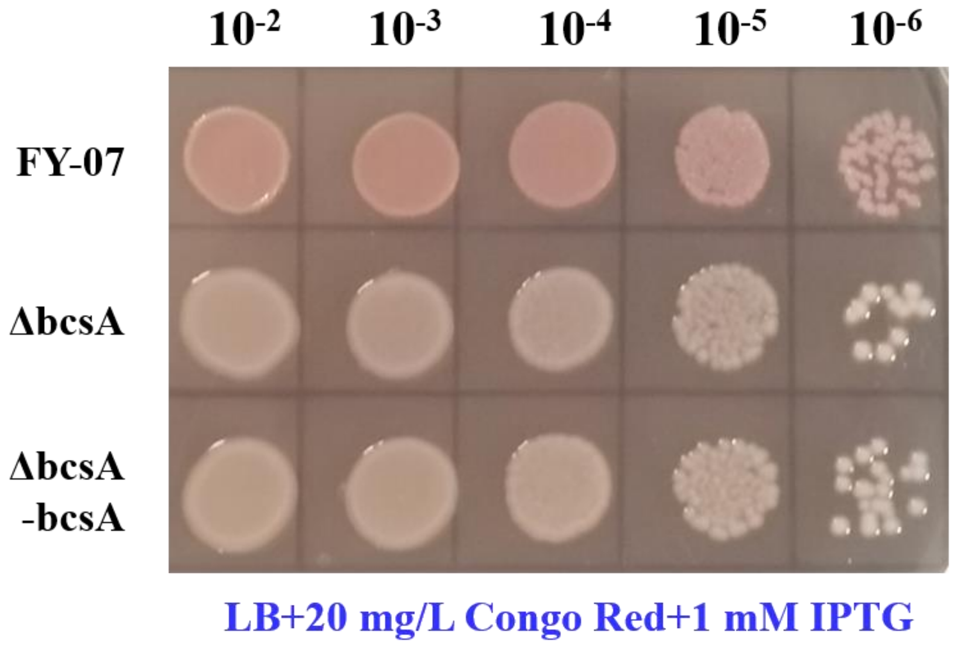


**Fig. S2 Comparison of the cellulose-synthesis ability of different strains**

**A**

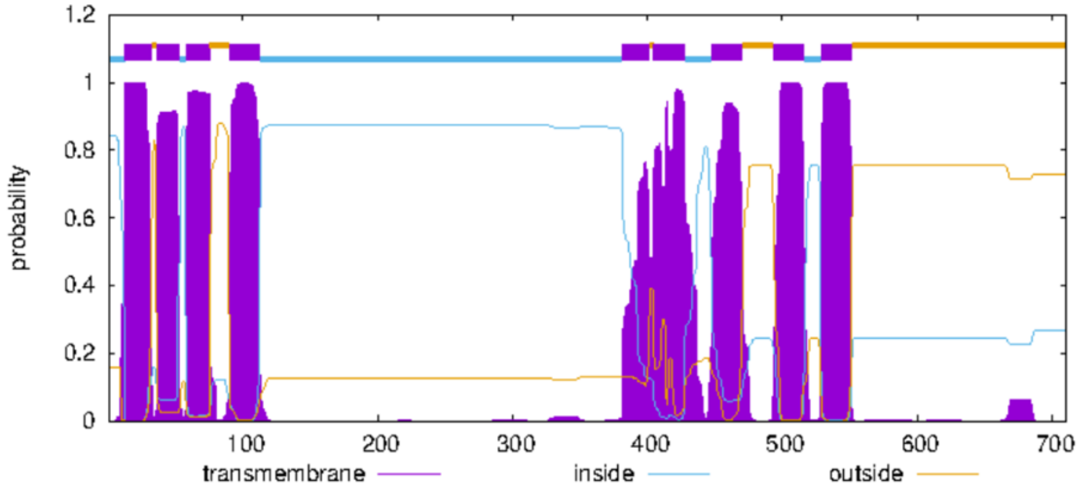


**B**

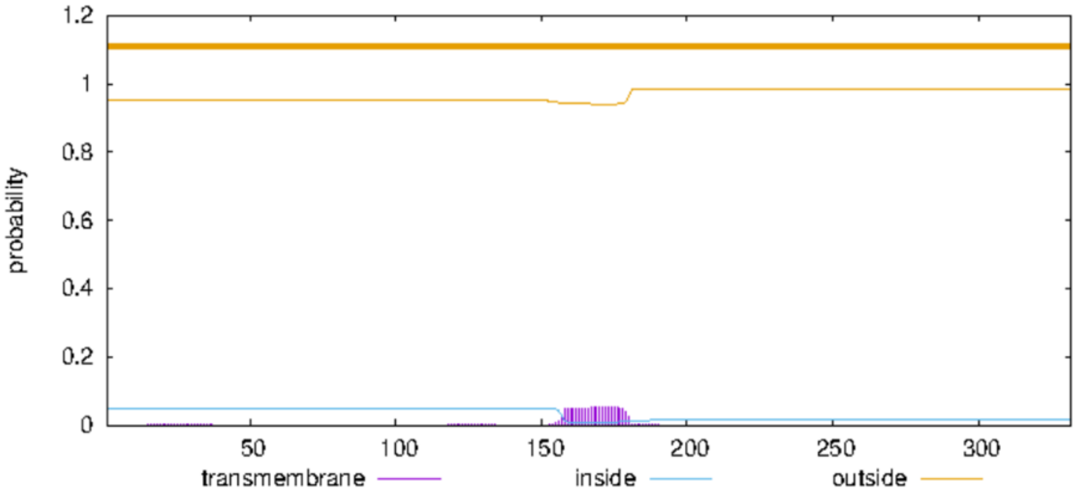


**Fig. S3 Transmembrane region prediction of BcsA and Fbp**

Note: A: Transmembrane region prediction of BcsA by an online tool (https://www.novopro.cn/tools/tmhmm.html), result indicated that BcsA contained eight transmembrane regions; B: Transmembrane region prediction of Fbp, result indicated that Fbp did not contain transmembrane region.

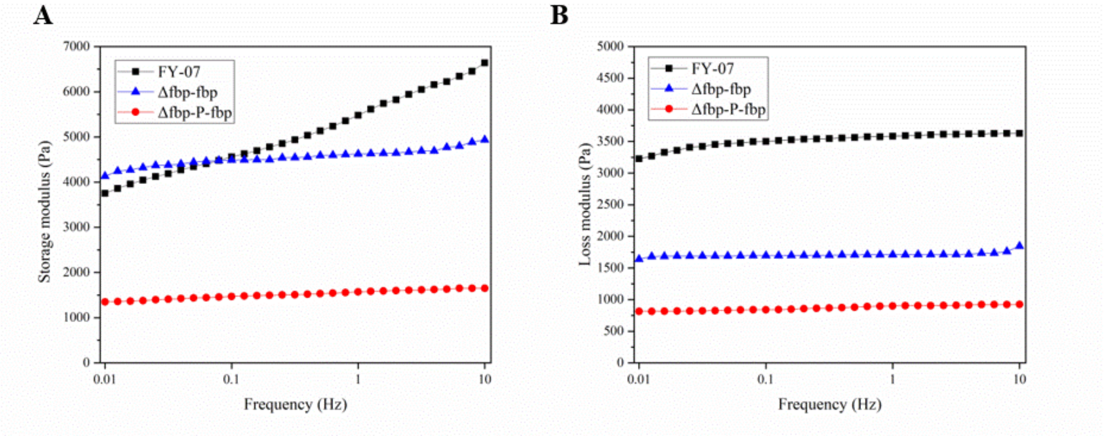


**Fig. S4 Rheological properties of BCs produced by *Enterobacter* sp. FY-07 and recombinant strains**

Note: A: Storage modulus (G'); B: Loss modulus (G").

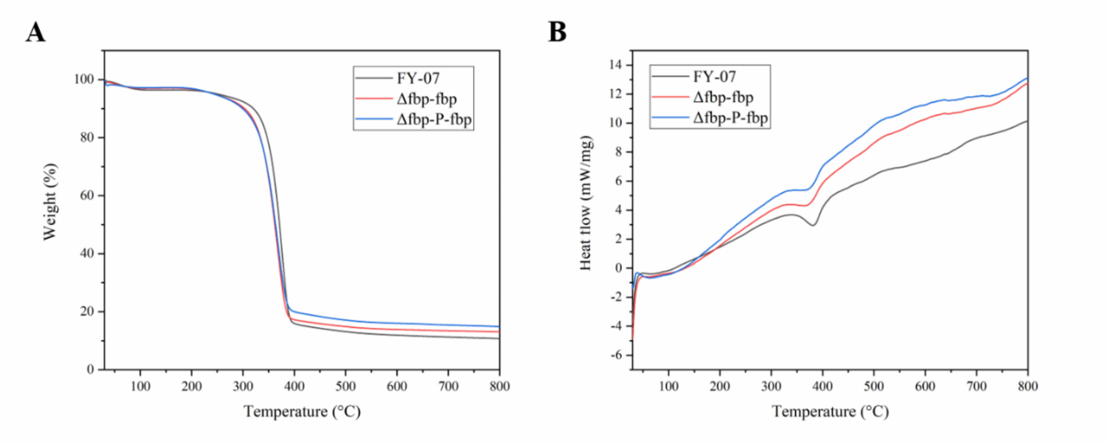


**Fig. S5 Thermogravimetric analysis (A) and differential scanning calorimetry (B) of BC produced by the wild-type and recombinant strains**

Table S1 Recombination efficiency of the λ Red-mediated gene knockout method.

| Genes name | Length of homologous arms (bp) | Number of colonies | Number of knocked-out strains | Recombination efficiency (%) |
|---|---|---|---|---|
| *bcsA* | 50 | 8 | 8 | 100 |
| *fbp* | 50 | 6 | 6 | 100 |

Table S2 Tensile test of BC pellicles produced by the wild-type and recombinant strains.

| Samples | Tension strength (MPa) | E% (%) | Young's modulus (MPa) |
|---|---|---|---|
| FY-07 | 7.254 ± 0.324 | 1.544 ± 0.071 | 0.524 ± 0.069 |
| Δfbp-fbp | 7.233 ± 0.815 | 2.214 ± 0.239 | 0.423 ± 0.095 |
| Δfbp-P-fbp | 6.586 ± 0.513 | 3.017 ± 0.194 | 0.307 ± 0.067 |
